# An *in silico* approach to determine inter-subunit affinities in human septin complexes

**DOI:** 10.1101/2022.12.20.521017

**Authors:** Benjamin Grupp, Justin A. Lemkul, Thomas Gronemeyer

**Affiliations:** Institute of Molecular Genetics and Cell Biology, James Franck Ring N27, Ulm University, 89081 Ulm, Germany; Department of Biochemistry, Virginia Tech, Blacksburg, VA 24061, USA

## Abstract

The septins are a conserved family of filament-forming guanine nucleotide binding proteins, often named the fourth component of the cytoskeleton. Correctly assembled septin structures are required for essential intracellular processes such as cytokinesis, vesicular transport, polarity establishment, and cellular adhesion. Structurally, septins belong to the P-Loop NTPases but they do not mediate signals to effectors through GTP binding and hydrolysis. GTP binding and hydrolysis are believed to contribute to septin complex integrity, but biochemical approaches addressing this topic are hampered by the stability of septin complexes after recombinant expression and the lack of nucleotide-depleted complexes. To overcome this limitation, we used a molecular dynamics-based approach to determine inter-subunit binding free energies in available human septin dimer structures and in their apo forms, which we generated *in silico*. The nucleotide in the GTPase active subunits SEPT2 and SEPT7, but not in SEPT6, was identified as a stabilizing element in the G interface as it is coordinated at its ribose ring to conserved amino acids. Removal of GDP from SEPT2 and SEPT7 results in flipping of a conserved Arg residue and disruption of an extensive hydrogen bond network in the septin unique element, concomitant with a decreased inter-subunit affinity.

## Introduction

Septins were discovered in budding yeast as cytoskeletal proteins and have now been identified in all eukaryotes except higher plants. Structurally, they are highly conserved, though the number of genes per organism is variable (e.g., 7 in *S. cerevisiae*, 13 in *H. sapiens* and *M. musculus*, 2 in *C. elegans*, and 1 in Chlamydomonas). Based on phylogenetic relationships, the septins can be sorted into five orthologous groups and members within one group can replace each other in higher ordered septin structures (Shuman and Momany 2022).

When the first septin crystal structure became available, it was confirmed that septins are guanine nucleotide binding proteins belonging to the superclass of P-loop NTPases, which was previously predicted by sequence analysis and shown by different *in vitro* assays (Farkasovsky et al. 2005; Sirajuddin et al. 2007; Valadares et al. 2017; Versele and Thorner 2004; Vrabioiu et al. 2004). The guanine nucleotide binding domains (G-domains) of the septins assemble into filamentous structures. The building block of these filaments in mammals is a linear, apolar, hexameric or octameric rod composed of the subunits SEPT2-6-7-(9)-(9)-7-6-2 (Mendonça et al. 2019, 2021; Sirajuddin et al. 2007). Septin filaments can be arranged into higher ordered structures like rings, cages, and gauzes *in vivo* and *in vitro* and have been shown to interact with actin and microtubules (Mostowy and Cossart 2012). They play important roles in chromosome segregation, cytokinesis, exocytosis, apoptosis, and also in the establishment of several diseases including cancer (Kinoshita 2003; Menon 2018; Mostowy and Cossart 2012; Peterson and Petty 2010; Weirich, Erzberger, and Barral 2008).

The polymerization into rods has been shown to be an important prerequisite for association with some binding partners as new topologies are formed along the subunit junctions (Sheffield et al. 2003). Septin rods exhibit a cyclic symmetry (C2) around the homotypic interface formed between the central rod subunits SEPT7 (in hexameric rods) or SEPT9 (in octameric rods). This symmetry seems to be maintained in all protofilament species formed as no asymmetrical incorporation of subunits has been observed. The inter-subunit contacts within the rods can be characterized as two distinct interfaces that appear in an alternating fashion: the G interface is solely stabilized by interactions of G domain components, whereas the NC interface is maintained by interactions between the N- and C-termini of two neighboring subunits. In the human canonical rod, the interface order is _NC_-SEPT2-_G_-SEPT6-_NC_-SEPT7-_G_-SEPT9-_NC_. Consequently, the central, homotypic interface in octameric rods is NC whereas it is G in hexameric rods lacking SEPT9. The availability of the structures of a trimeric and a hexameric rod allowed for the identification of all interface components (reviewed in (Cavini et al. 2021)). G-interfaces are stabilized by interactions between the P-loop, switch 1, switch 2, G4, trans loops 1 and 2, and the bound nucleotides. Hydrophobic interactions between the septin unique elementβ -meander (SUE-βββ) of both subunits contribute additionally to interface stability (Fig. 1A). Mutation of a conserved Trp maintaining these interactions disrupt the interface (Sirajuddin et al. 2007).

**Figure 1.**
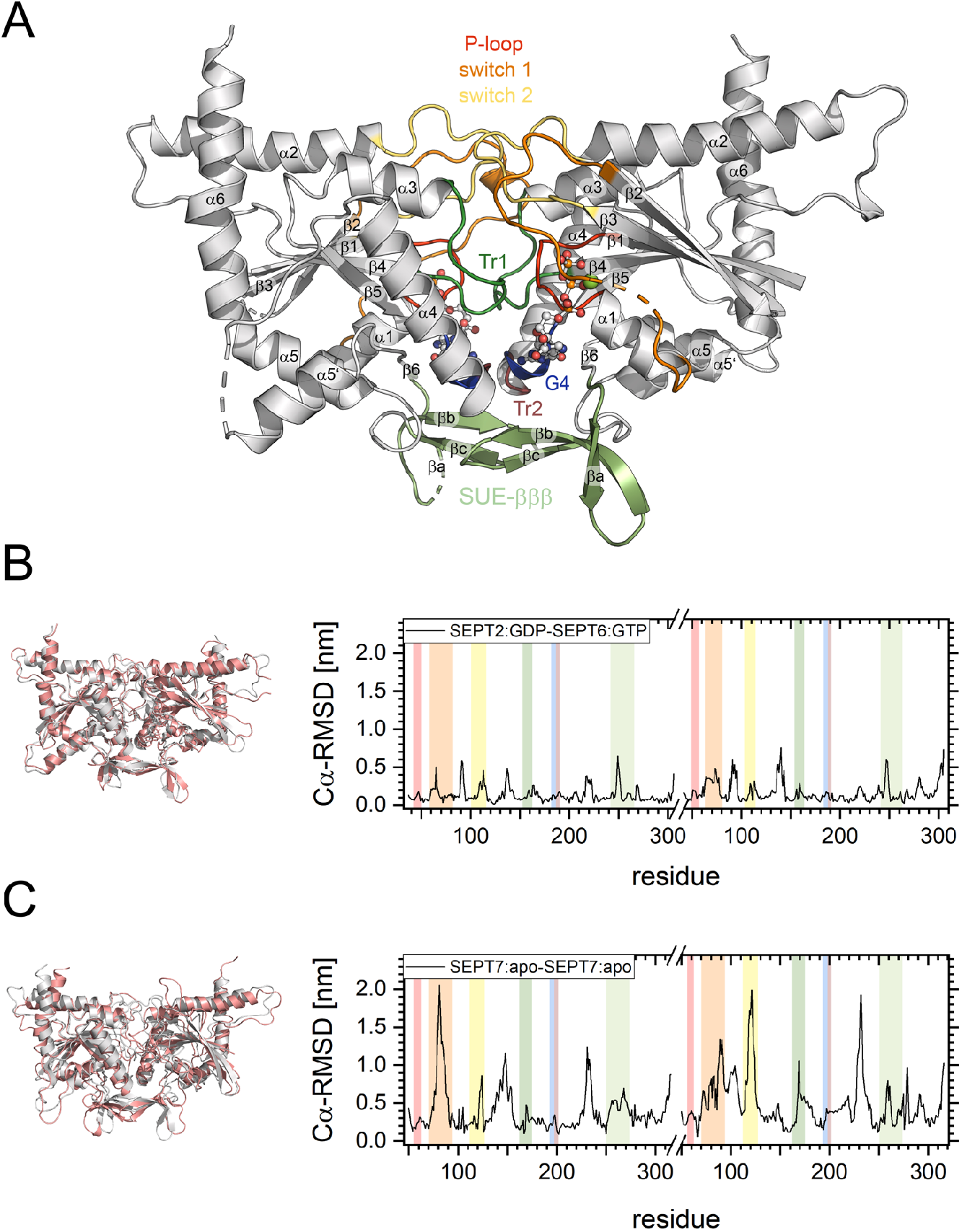
Overview of the employed molecules.**(A)** G interface between the human septins SEPT2:GDP (left) and SEPT6:GTP (right) (PDB-ID 6UPA). Sheets and helices are numbered according to the classical G domain numbering of Ras-like proteins. Features of the septin G domain are annotated and colored; SUE: septin unique element; Tr: trans-loop. The GDP in SEPT2 and the GTP and the Mg^2+^ in SEPT6 are shown as ball-and-stick presentation. **(B)** Cα-RMSD of the relaxed structures referenced to the input structures of SEPT2:GDP-SEPT6:GTP (least deformed structure) and **(C)** SEPT7:apo-SEPT7:apo (most deformed structure). An overlay of the relaxed structure (salmon) with the input structure (grey) (left) and a RMSD plot (right) is shown. The color coding in the RMSD plots corresponds to the structural elements highlighted in (A)

Much less is known about the process and the kinetics of protofilament formation and the influence of the nucleotide on this process. This question was, until now, only addressed in yeast. In 2017, Weems et al. suggested a model for the step-wise pathway of septin protofilament assembly in *S. cerevisiae* based on data obtained from an *in vivo* Bimolecular Fluorescence Complementation assay. In addition to revealing the order of septin interactions that are established during protofilament assembly, they predicted active communication between G- and NC-interfaces mediated by either the nucleotide binding state or the association of the native septin binding partner (Weems and McMurray 2017). Comparable information regarding human septins is missing due to the complexity of the corresponding *in vivo* experimentation. Recombinantly expressed mammalian septin subunits usually form stable dimers (Brognara et al. 2019; Castro et al. 2020; Macedo et al. 2013; Rosa et al. 2020; Sirajuddin et al. 2007; Zent, Vetter, and Wittinghofer 2011) with high affinity between the protomers, hampering direct detection of intra-complex affinities. However, the dissociation of SEPT7 mutants with lowered affinity was shown to be dependent in the nature of the nucleotide (Zent and Wittinghofer 2014).

The availability of supercomputers and high-performance clusters has enabled the use of *in silico* approaches aiming at modelling processes that cannot be accessed in wet-lab experiments. Specifically, molecular dynamics (MD) simulations produce biophysically meaningful trajectories that reveal not only thermodynamic but also dynamic information (Klepeis et al. 2009; Zheng et al. 2018). The dynamics of a given system are propagated via an equation of motion that describes how atomic coordinates evolve in fixed time increments. Dynamics of binding-unbinding events in proteins, such as protein domain unfolding or antibody-antigen binding, are often studied by steered molecular dynamics simulations (SMD) (reviewed in (Isralewitz et al. 2001)). In SMD, external forces (often a harmonic biasing potential) are applied to a system (e.g., a protein complex) to probe mechanical functions and to accelerate processes which are otherwise too slow to be observed within timeframes accessible to typical unbiased MD simulations. By “pulling” the different components apart from each other along a selected reaction coordinate, events such as protein/ligand or protein/protein unbinding can be studied.

Using a biasing potential to drive such a system from one state to another allows for the calculation of free energy changes by a method called umbrella sampling (US) (reviewed in (Kästner 2011)). US is usually performed in a series of individually modelled sampling windows, each representing a position along the reaction coordinate between the end states. These windows can be debiased with e.g. the weighted histogram analysis method (Kumar et al. 1992) to calculate the potential of mean force (PMF) and eventually protein-protein binding free energies (Patel and Ytreberg 2018). US has been applied to a number of biological problems inaccessible to wet-lab experimentation such as to assess the stability of amyloid protofibrils (Lemkul and Bevan 2010) or switch peptide binding to cardiac troponin (Cool and Lindert 2022), to name only two examples.

We present here the application of US based MD simulations to investigate the contribution of bound nucleotide to binding affinities between septin subunits in human septin dimers making use of available structural data.

## Methods

### Preparation of coordinate files

The human septin dimers SEPT2-SEPT6 (PDB-ID 6UPA) and SEPT7-SEPT7 (PDB-ID 6N0B) were used for performing the MD simulations. Unresolved regions were modeled in using the SWISS-MODEL User Template Mode. N- and C-terminal extensions were cropped at sequentially related sites, leading eventually to the dimers [SEPT2_E37-R306_:GDP] – [SEPT6_C42-E305_:GTP:Mg] and [SEPT7_E50-A316_:GDP] – [SEPT7_E50-A316_:GDP].

Each dimer structure was further processed using Maestro (Schrödinger Release 2021-1, Maestro, Schrödinger, LLC, New York, 2021). Assignment of the protonation state was performed at pH 7, followed by optimization of the hydrogen bond networks via 180° flipping of terminal amide groups in Asn/Gln and the ring of His. The structures were then relaxed using the OPLS4-ff (Lu et al. 2021) until either convergence of the potential energy or a heavy atom RMSD of 0.30 Å was reached (default upper limit RMSD in Maestro) (Madhavi Sastry et al. 2013). This was followed by another round of hydrogen bond optimization.

The residue and atom names in the output Maestro PDB file were made compliant with the CHARMM36m force field using the CHARMM-GUI webserver (Huang et al. 2017; Lee et al. 2016). The N- and C-termini of each structure were capped in PyMOL with acetyl and N-methyl amide groups, respectively, to mimic the full-length, uncharged state at these positions. For simulations using the apoproteins, the bound ligands were deleted at this point.

### Unbiased simulations

All simulations described were performed using GROMACS v. 2022.2 (Abraham et al. 2015) using the CHARMM36m force field (Huang et al. 2017) on the JUSTUS2 high performance cluster at Ulm University. Bonds with hydrogen atoms were constrained using the LINCS algorithm (Hess 2008; Hess et al. 1997), allowing the time step for the integration to be set to 2 fs. Short-range Lennard-Jones interactions were smoothly switched to zero between 1.0 - 1.2 nm. Electrostatic interactions were calculated using the smooth particle mesh Ewald (PME) algorithm (Darden, York, and Pedersen 1998; Essmann et al. 1998) with a real-space cutoff of 1.2 nm. No dispersion correction was applied. Periodic boundary conditions were applied in all directions.

For unbiased MD simulations, each dimer was placed into a dodecahedral box with dimensions chosen to ensure a minimum distance of 3.0 nm to the periodic image in the initial conformation. The box was filled with water molecules described by the CHARMM-modified TIP3P model (Durell, Brooks, and Ben-Naim 1994; Jorgensen et al. 1998; Neria, Fischer, and Karplus 1998). A total of 150 mM NaCl was added to the system, including neutralizing counterions. Each system was relaxed via steepest descent minimization with a target maximum force of 500 kJ·mol^-1^·nm^-1^. Next, the system was equilibrated to a target temperature of 310 K over 50 ps under an NVT ensemble using the velocity rescaling thermostat (Bussi, Donadio, and Parrinello 2007) to regulate temperature with a time constant of 0.1 ps. Position restraints of 1000 kJ·mol^-1^·nm^-2^ were applied to heavy atoms in the proteins and associated ligands. Initial velocities were assigned randomly according to the Maxwell-Boltzmann distribution. Equilibration was continued under an NPT ensemble to allow the systems to reach the target pressure (1.0 bar) over 500 ps. Pressure was regulated isotropically with the stochastic cell rescaling barostat (Bernetti and Bussi 2020) and a time constant of 2 ps was used for the pressure coupling. The same position restraints were maintained during this phase of equilibration.

Production simulations were conducted under the same conditions as the NPT equilibration but without restraints and with pressure regulated by the Parrinello-Rahman barostat (Parrinello and Rahman 1998). Three independent production simulations (500 ns) were performed for septin dimer relaxation before COM-pulling simulations.

### COM-pulling and umbrella sampling (US) simulations

Input structures for COM-pulling simulations were generated by pooling the last 200 ns of the three individual unbiased simulations, followed by backbone RMSD-based clustering via the algorithm published by Daura et al. (Daura et al. 1998) after least-squares fitting. The cutoff was set to 2 Ả with 100 ps increments between sampled states. The central structure in the largest cluster was then used in the following simulations. The septin dimers were then oriented such that the filament axis coincided with the x-axis of a rectangular box with the x/y/z dimensions 22/10/10 nm. The box was subsequently solvated, minimized, and equilibrated as described above. Steered simulations were subsequently performed without position restraints. Different combinations of pull rate v (0.001 – 0.05 nm·ps^-1^) and spring constant k (100 – 3000 kJ·mol^-1^·nm^-2^) were analyzed regarding low structural deformation and reproducible maximum forces, yielding eventually a spring constant of 500 kJ·mol^-1^·nm^-2^ and a pull rate of 0.0075 nm·ps^-1^ that were used in the COM-pulling simulations. Both dimer subunits were separated until a COM-distance (as defined by the Cα atoms of the proteins) of approximately 10 nm was reached.

For US, different configurations were extracted from the COM-pulling simulations with a symmetric window spacing of ∼0.15 nm and a COM distance until ∼7.0 nm. Windows were selected using the scripts described in the GROMACS umbrella sampling tutorial (Lemkul 2019). Higher COM distances were not considered since the slope of the potential of mean force (PMF) curves asymptotically approached zero within COM distances below 7.0 nm. The extracted frames were equilibrated via a 500 ps NPT simulation to the target temperature (310 K) and target pressure (1.0 bar) using the same restraints described above. In the following 20 ns production simulations (under an NPT ensemble), no restraints were applied, and the Cα-COM separation of the subunits in each window was maintained via a harmonic potential along the x-axis defined by a spring constant of 1000 kJ·mol^-1^·nm^-2^. When necessary, additional simulations were performed for unsampled regions using a smaller window spacing of 0.05 nm and a spring constant of 2500 kJ·mol^-1^·nm^-2^. PMF profiles were then generated using the WHAM method (Kumar et al. 1992) based on the Cα-COM separations observed in the different windows between 5 – 20 ns of the simulations (Hub, De Groot, and Van Der Spoel 2010). ΔG_bind_ was determined by subtracting the highest energy value from the lowest one observed in the PMF. COM-pulling simulations and US were performed in quintuplicate for each construct.

## Results and Discussion

The impact of nucleotide binding and hydrolysis on septin protofilament formation and the polymerization properties of protofilaments into higher ordered structures remains largely enigmatic. Previous research indicated that the presence of nucleotide has a promoting effect on yeast septin filament formation *in vitro* on lipid bilayers (Bertin et al. 2010). Recombinantly expressed individual septin subunits form either stable dimers or even multimers and all crystal structures of G interface dimers available in the PDB contain nucleotides at the interface. Only one available crystal structure (of the yeast septin Cdc11, PDB-ID 5AR1) shows a monomer in the nucleotide-free apo form and exhibits an unusually positioned SUE-βββ(Brausemann et al. 2016). Together, these findings suggest that the nucleotide plays a pivotal role in the formation of G interfaces within septin complexes. Knowledge of the inter-subunit affinities within a septin complex in the nucleotide-bound and apo states would allow for conclusions to be drawn regarding the stability of the complexes and the dependence on the nucleotide. Since direct measurement of affinities *in vitro* is hindered by the stability of the protomers and dimers lacking nucleotides have never been generated experimentally, we established a molecular dynamics (MD) simulation pipeline based on available structural data to address this issue.

We selected the crystal structures of the SEPT2_GDP_-SEPT6_GTP:Mg_ (PDB-ID 6UPA) (Rosa et al. 2020) and SEPT7_GDP_-SEPT7_GDP_ (PDB-ID 6N0B) (Brognara et al. 2019) G interface dimers for investigation. These dimer structures represent the highest-resolution structures available for the G interface partners in the canonical human hetero-hexameric septin protofilament complex (Mendonça et al. 2021). Consistent with this hexameric structure, only SEPT6 contains GTP, while SEPT2 and SEPT7 contain GDP. Missing and/or unresolved regions in the structures were completed *in silico* and we subsequently removed one or both nucleotides from each dimer, resulting in the nucleotide:nucleotide, nucleotide:apo, apo:nucleotide and apo:apo states. Each generated dimer structure was relaxed in unrestrained simulations to obtain equilibrated starting structures for the COM-pulling simulations. These structures are shown in Suppl. Fig. 1. Overall, the structural integrity of all employed septin pairs was maintained, with SEPT7:apo-SEPT7:apo showing the highest degree of deformation compared with the unrelaxed input structure. The Cα-RMSD of the relaxed structures referenced to the input structures of the least- and most-deformed septin dimer (SEPT2:GDP-SEPT6:GTP and SEPT7:apo-SEPT7:apo, respectively) are illustrated in Fig. 1B, C (see Table 1 and Suppl. Fig. 2 for all RMSD values and RMSD plots).

**Table 1.** Cα-RMSD values of the central structures of the largest clusters after unbiased MD simulations compared with the initial structures.

| system | Cα-RMSD [nm] |
| --- | --- |
| **SEPT2 - SEPT6** |  |
| SEPT2:GDP - SEPT6:GTP | 0.191 |
| SEPT2:GDP - SEPT6:apo | 0.230 |
| SEPT2:apo - SEPT6:GTP | 0.224 |
| SEPT2:apo - SEPT6:apo | 0.223 |
| **SEPT7 - SEPT7** |  |
| SEPT7:GDP - SEPT7:GDP | 0.331 |
| SEPT7:GDP - SEPT7:apo | 0.299 |
| SEPT7:apo - SEPT7:GDP | 0.328 |
| SEPT7:apo - SEPT7:apo | 0.536 |

Closer inspection of the deformed regions reveals a pivotal influence of the nucleotide on G interface stability. The highly conserved Arg(βb), Tyr(βb) and Asp(G4) (following the recently established nomenclature “residue(motif)” (Cavini et al. 2021)) coordinate the ribose ring of the nucleotide by multiple hydrogen bonds (Fig. 2). Arg(βb) is itself coordinated by the Asp(G4) and an equally conserved Glu(Tr2) from the neighboring subunit. Removal of the GDP from SEPT2 and SEPT7 results in a 90° flipping of Arg(βb). This flipping abolishes the coordination with Asp(G4), disrupts the hydrogen bond network and might be responsible for the conformational deformation in the SUE-βββ(Fig. 2A, B). In SEPT6, the positioning of Arg(βb) and its coordination with Asp(G4) and Glu(Tr2) is fully maintained after removal of the GTP (Fig. 2C). Consequently, the conformational deformation is highest in the SEPT7:apo-SEPT7:apo dimer since two Arg(βb) flipping events destabilize the G interface.

**Figure 2.**
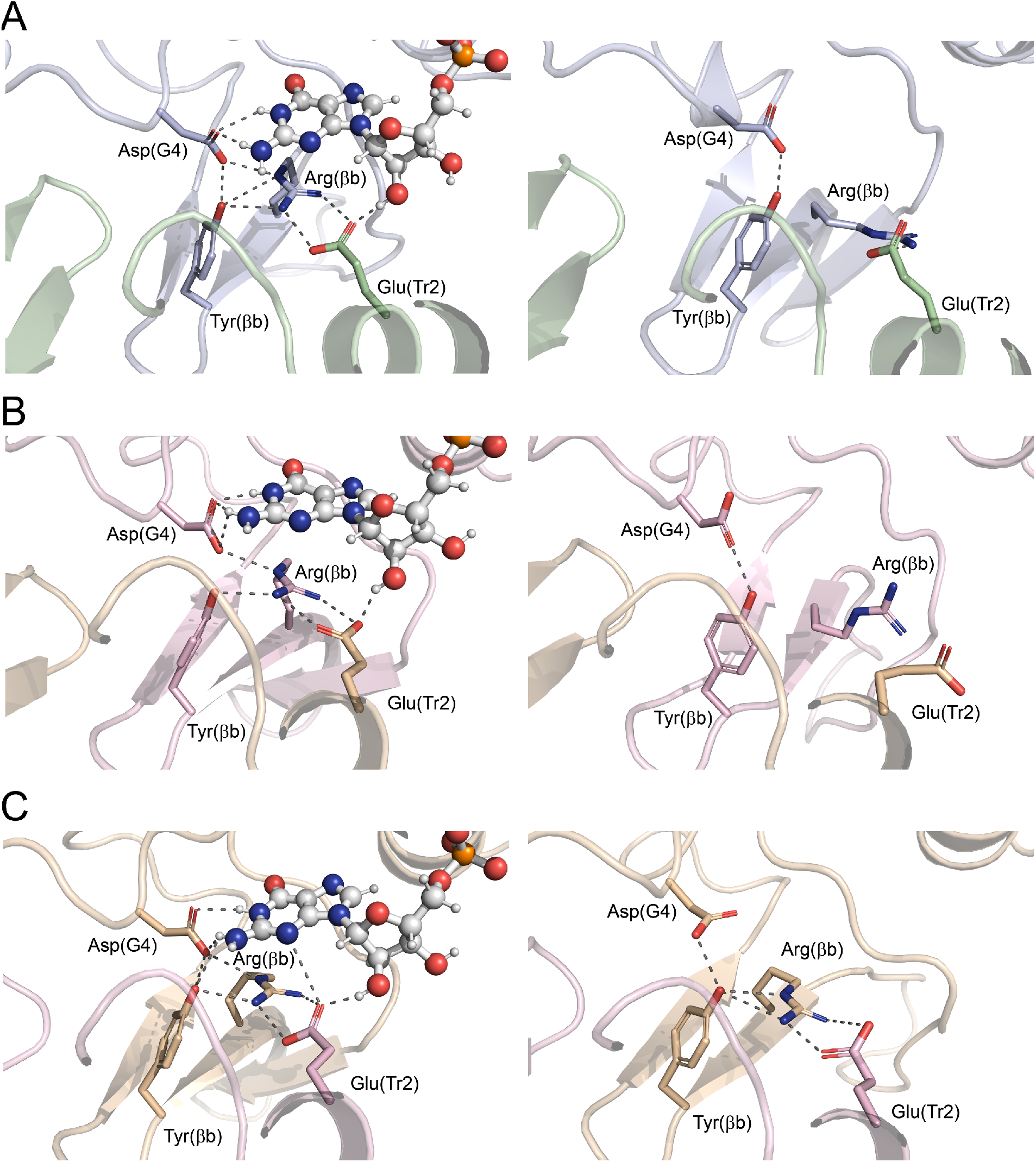
Structural rearrangement in the binding pocket after nucleotide removal. Arg(b) flipping upon nucleotide removal in **(A)** SEPT7, **(B)** SEPT2 and **(C)** SEPT6. Note that the Arg(b) in SEPT6 remains in its position and that no flipping occurs after GTP removal. Shown is a detail of the G interface in the nucleotide bound state (left) and in the apo state (right). Arg(βb), Tyr(βb), Asp(G4) and Glu(Tr2) are shown in stick presentation and the nucleotide is shown in ball and stick presentation, respectively. SEPT7 subunits are colored palegreen and lightblue, SEPT2 lightpink and SEPT6 wheat.

The linear arrangement of septin subunits along a (proto-)filament axis allowed us to place the dimers in a rectangular box with the filament axis aligned with the x-axis for the following COM-pulling simulations (Fig. 3A). We tested the influence of different spring constants and pulling velocities on the system. The spring constant, k, determines the stiffness of the one-dimensional harmonic potential that is applied to increase the COM distance of the two septin subunits with the pulling velocity, v. Average values for k (500 kJ·mol^-1^·nm^-2^) and v (0.0075 nm·ps^-1^) resulted in reproducible simulation runs with a low standard deviation of the dissociation force and low structural deformation (Suppl. Fig. 3). COM-pulling of the Cα atoms in each subunit was then performed along the x-axis, assuming a one-dimensional reaction coordinate based on the filamentous structure of septin (proto-)filaments. A typical dissociation pathway of two subunits within a septin dimer (here, SEPT2:GDP-SEPT6:GTP) is shown in Fig. 3A-D. Reaching the maximum applied force, the two subunits begin to dissociate (Fig. 3B, corresponding to the blue dot in the force vs. time plot in Fig. 3E). Structural transitions after full separation of the dimers (Fig. 3 C and D, with each state indicated in Fig. 3E) are minor, although the G interface part becomes solvent-exposed.

**Figure 3.**
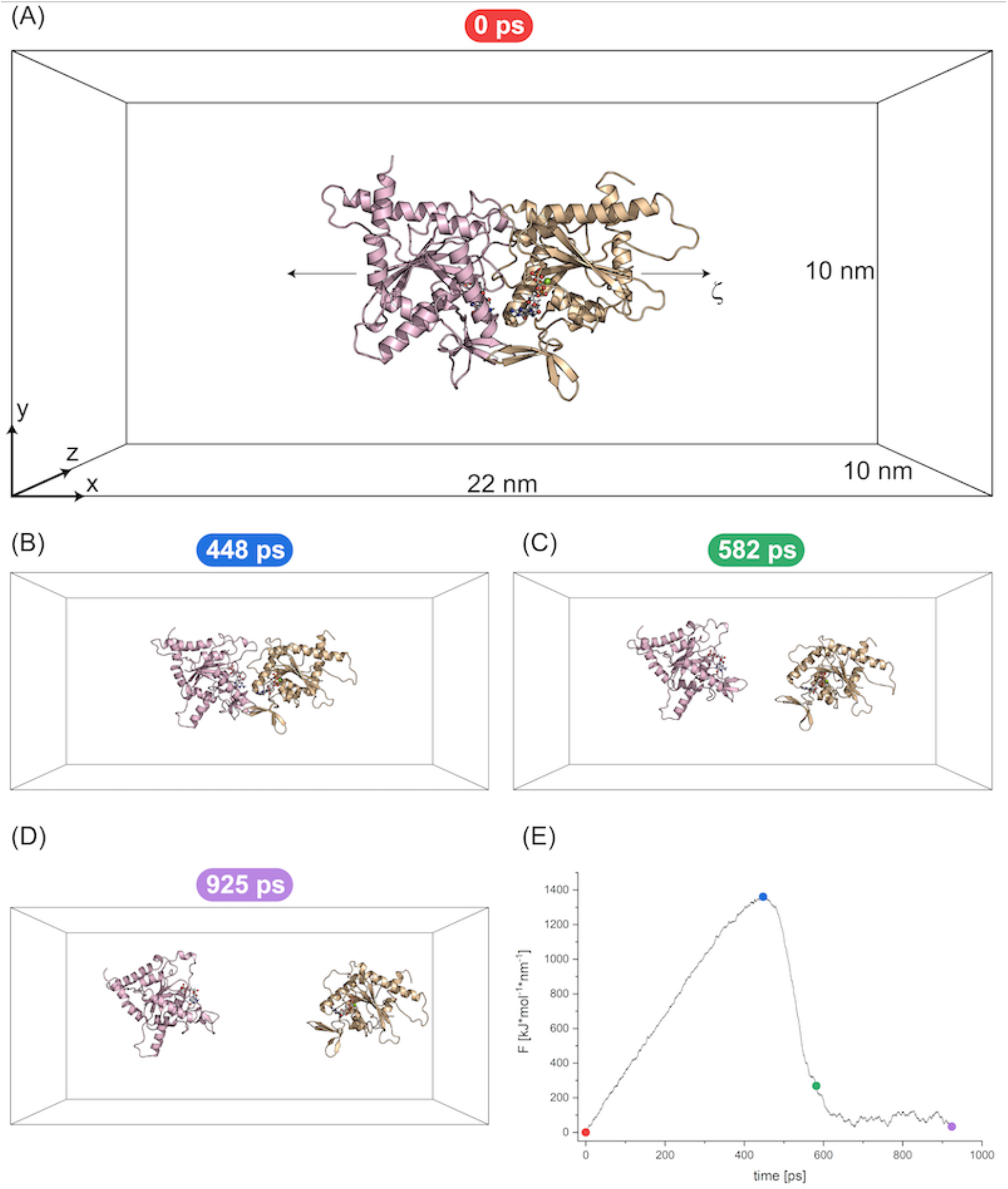
Representative COM-pulling simulation. **(A)** SEPT2:GDP-SEPT6:GTP dimer inside the simulation box before force application. The box dimensions, axis orientation, and the reaction coordinate ζ are indicated. Note that ζ corresponds to the x-axis. **(B)-(D)** selected time points of the trajectory with **(B)** maximum force application (dissociation point), **(C)** last timepoint of the simulation where intramolecular interactions may occur; i.e., protein atoms are in distance of 1.2 nm, **(D)** endpoint of the simulation.**(E)** Force vs. time plot of the SEPT2:GDP-SEPT6:GTP simulation. The timepoints shown in A-D are indicated as dots.

Comparing the trajectories of the SEPT2-SEPT6 and the SEPT7-SEPT7 dimers reveals a striking difference. Pulling SEPT2 and SEPT6 apart resulted in an overall linear movement along the reaction coordinate with a modest tilt (Video 1). The entangled SUE-βββslide versus each other until the maximum force is reached. The SEPT7 subunits, however, are separated in a strongly tilted movement with the pivotal point at the SUE-βββ. The upper part of the interface bearing Tr1 and the switch 1 and 2 regions (Fig. 1A) is already fully separated while the SUE-βββstill interact (Video 2). Tilting continues even after full separation of the two subunits. To quantify this phenomenon, we determined the tilt angle between two subunits during a COM-pulling simulation. The α2 helix is oriented almost parallel to the filament axis (Fig. 1A) and is therefore best suited as a reference structure element to quantify tilting between two adjacent subunits relative to the reaction coordinate. We defined a linear helical axis in α2 of each subunit in a dimer and recorded the angle between the two axes over the whole trajectory (Fig. 4). The tilt angle becomes distinctly larger in all SEPT7-SEPT7 separations compared to SEPT2-SEPT6 separations. However, whereas all tested SEPT7-SEPT7 states show the same prominent degree of tilting, the SEPT2-SEPT6 dimers are more variable, with some states adopting more tilted orientations than others (reflected by the larger standard deviation shown in Fig. 4A).

**Figure 4.**
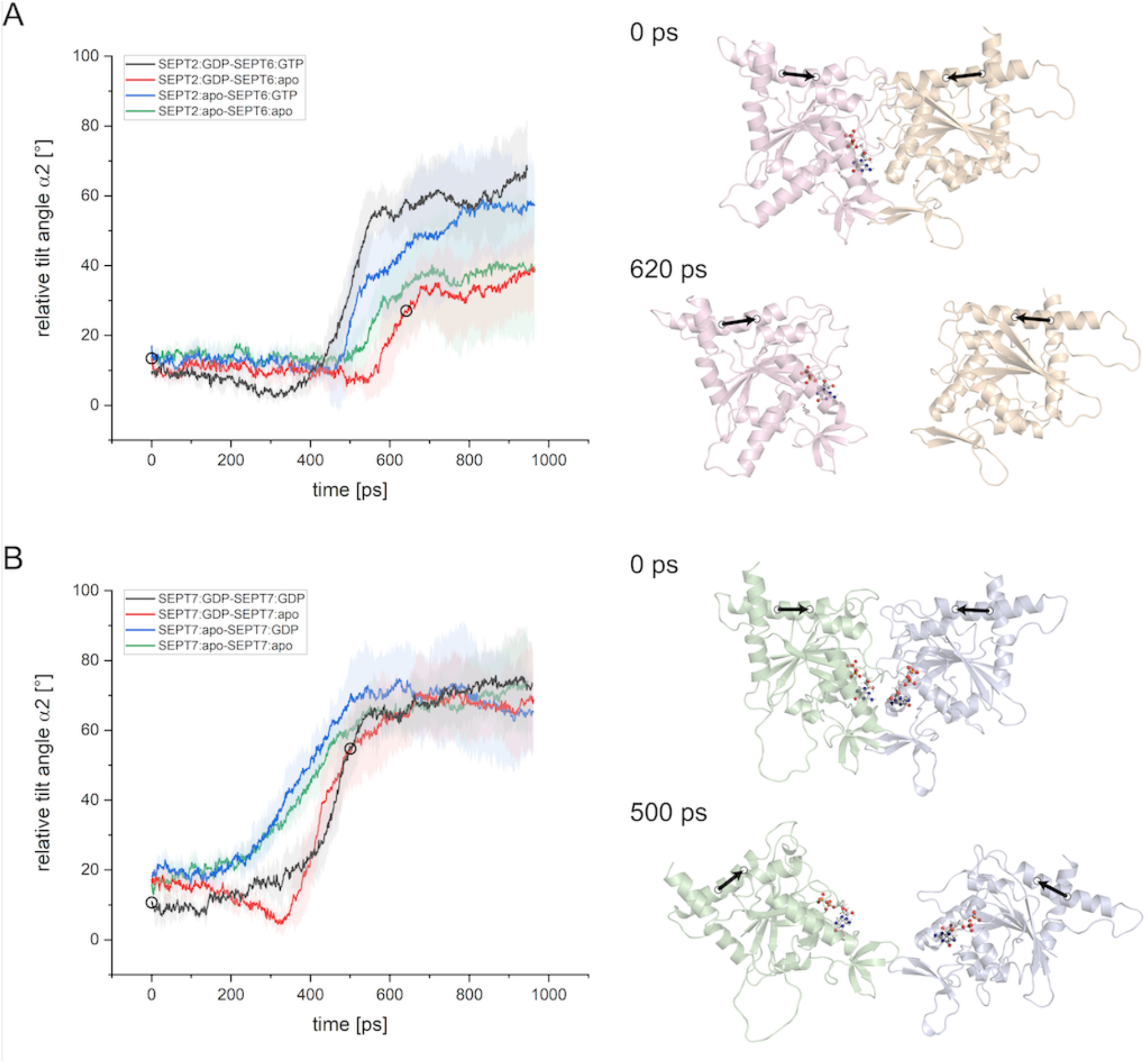
Tilting between septin subunits during COM-pulling simulations. The tilt angle was plotted vs. the simulation time (left) for **(A)** SEPT2-SEPT6 and **(B)** SEPT7-SEPT7 dimers. Curves represent average curves of five independent trajectories, error bands represent the standard deviation. Two conformations, one at the starting point and one shortly before reaching the maximum tilt angle (indicated as circles in the plots), are shown (right), demonstrating enhanced tilting in SEPT7 dimers. COM points defining the central helical axis (shown as an arrow) in the α2 helix are indicated.

The tilt angle vs. time plots (Fig.4) do not reveal a prominent influence of the nucleotide on tilting. Since the SUE-βββ sequence is highly conserved among septins (Cavini et al. 2021), we propose that the interactions between the switch- and Tr1 loops in the upper part of the interface are weaker in the SEPT7 dimers than in the SEPT2-SEPT6 dimers, promoting a disruption of this region before dissociation of the SUE-βββtakes place.

Subsequently, the binding free energies between two septin subunits within a dimer were determined via US simulations. Therefore, different conformations along the reaction coordinate ζ were extracted from the corresponding COM-pulling simulations for Cα-COM-separation distances between approx. 3.5 nm (representing the starting conformation) and approx. 7.0 nm. The resulting one-dimensional PMF profiles for the SEPT2-SEPT6 and SEPT7-SEPT7 dimers are shown in Fig. 5A and 5B, respectively, allowing for the calculation of ΔG via the weighted histogram analysis method (WHAM) (Hub et al. 2010; Kumar et al. 1992). Fig. 5C shows an example for the Cα-COM-separation distribution in one umbrella window. All obtained ΔG values are plotted in Fig. 5D and listed in Table 2.

**Table 2.** Binding free energies for the assessed septin dimer configurations

| system | $\Delta G$ [kJ·mol<sup>-1</sup>] | $\Delta\Delta G$ [kJ·mol<sup>-1</sup>] |
| --- | --- | --- |
| **SEPT2 - SEPT6** |  |  |
| SEPT2:GDP - SEPT6:GTP | -114 ± 10 | - |
| SEPT2:GDP - SEPT6:apo | -129 ± 7 | -15 ± 13 |
| SEPT2:apo - SEPT6:GTP | -95 ± 13 | 18 ± 17 |
| SEPT2:apo - SEPT6:apo | -95 ± 7 | 18 ± 13 |
| **SEPT7 - SEPT7** |  |  |
| SEPT7:GDP - SEPT7:GDP | -144 ± 11 | - |
| SEPT7:GDP - SEPT7:apo | -90 ± 6 | 55 ± 13 |
| SEPT7:apo - SEPT7:GDP | -105 ± 6 | 39 ± 13 |
| SEPT7:apo - SEPT7:apo | -63 ± 8 | 81 ± 14 |

**Figure 5.**
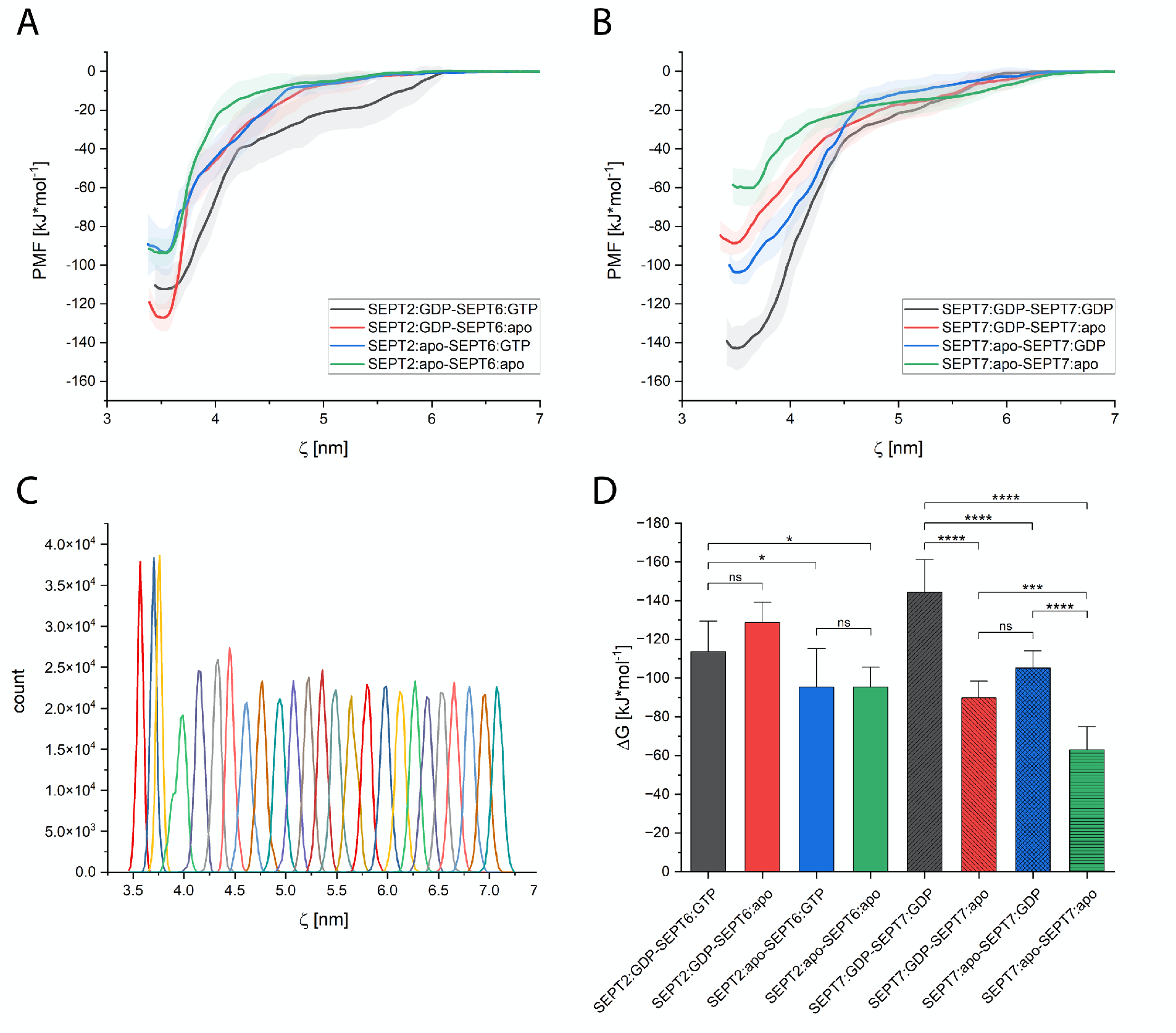
Determination of binding free energies using umbrella sampling. **(A)** PMF profiles for the SEPT2-SEPT6 dimers. **(B)** PMF profiles for the SEPT7-SEPT7 dimers. Error bands in (A) and (B) represent the standard deviation. **(C)** Representative umbrella sampling of SEPT2:GDP-SEPT6:GTP. Shown are the probability distributions of all sampled windows of one umbrella sampling series. **(D)** Graphical representation of the determined ΔG values (mean of five independent calculations) for each septin dimer. Error bars represent the standard deviation. P-values were determined via ANOVA followed by a Tukey posthoc analysis.

Removal of one or both nucleotide molecule(s) has a different impact on SEPT2-SEPT6 than on SEPT7-SEPT7. In SEPT7 dimers, the ΔΔG increases by (averaged) 46 kJ·mol^-1^ when one GDP is removed (difference between GDP-apo and apo-GDP not significant, see Fig. 5C). Removal of both nucleotides results in a significant increase by 81 kJ·mol^-1^ (Fig. 5C, Table 2). Consequently, both GDP molecules contribute to the binding affinity. In SEPT2-SEPT6, only the nucleotide in SEPT2 contributes to ΔG. While removal of the GTP from SEPT6 results in a 1′1′G differing insignificantly from the GDP-GTP state (Fig. 5C, Table 2), removal of the GDP from SEPT2 results in a ΔΔG-increase of approx. 18 kJ·mol^-1^, regardless of the presence of GTP in SEPT6 (Fig. 5C, Table 2).

What is the difference in the two complexes that might explain this phenomenon? SEPT2 and SEPT7 are both GTPase active whereas SEPT6 is not (reviewed in (Grupp and Gronemeyer 2022)). Consequently, our finding implicates that only the GDP from the two GTPase active subunits contributes to the binding affinity. This is in perfect agreement with the above described Arg(βb) flipping which occurs only in the GTPase active subunits upon nucleotide removal. We used the COM-pulling simulations to further substantiate the Arg(βb) flipping mechanism and the stabilizing role of the GDP by extracting the nonbonded energies for Arg(βb), Tyr(βb), Asp(G4) and Glu(Tr2) from all available states. Whereas the charged interactions remain unchanged in SEPT6, a pronounced weakening of the nonbonded energies can be detected in SEPT7 and SEPT2 when one or both GDP molecules are removed from one dimer (Suppl. Fig.4).

The human septin protofilament has the order *SEPT2-_G_-°SEPT6-_NC_-*SEPT7-_G_-*SEPT7-_NC_-°SEPT6-_G_-*SEPT2 (hexamer) or *SEPT2-_G_-°SEPT6-_NC_-*SEPT7-_G_-*SEPT9-_NC_-*SEPT9-_G_-*SEPT7-_NC_-°SEPT6-_G_-*SEPT2 (octamer, with GTPase active subunits marked with * and inactive subunits marked with °). SEPT7 forms always a G interface with another GTPase active septin (i.e., with another SEPT7 or with SEPT9) and SEPT2 forms always a G interface with the inactive SEPT6. It is thus tempting to speculate that GTP hydrolysis promotes G interface stability by providing GDP.

The role of nucleotide hydrolysis in septins is still poorly understood and subject of debate (Grupp and Gronemeyer 2022). GTP hydrolysis was shown by biochemical assays for members of the Momany 1a and 2b groups, but hydrolysis products are not released to the surrounding medium by native hexameric or octameric complexes (Fischer et al. 2022). It is meanwhile commonly accepted that septins evolved from a common ancestor (Pan, Malmberg, and Momany 2007) and during evolution some septins seem to have lost their ability to hydrolyze GTP. However, it can be anticipated that all septins are loaded with GTP during folding as GTP is the predominant intracellular guanine nucleotide. GTP hydrolysis may be the event that finalizes septin protofilament assembly by “locking” the G interface in its most stable (GDP bound) state. This model is in line with our finding that the SEPT7:GDP-SEPT7:GDP pair has an overall more favorable binding free energy than the SEPT2:GDP-SEPT6:GTP pair (Fig. 5C). Such a singular “lock-hydrolysis” mechanism would also explain the conundrum of hydrolysis products that can only be detected in denatured but not in native complexes. A singular hydrolysis event does not necessitate the release of hydrolysis products and further uptake of fresh GTP for another round of hydrolysis.

Zent and Wittinghofer determined binding affinities experimentally in SEPT7 dimers bound either to GDP or the non-hydrolyzable GTP analogon GppNHp by introducing affinity-lowering mutations in the G interface. Without mutations, the dimers did not dissociate at all. The dissociation rates in the mutants were 120-fold faster in the GppNHp than in the GDP state (Zent and Wittinghofer 2014), which corroborates our finding that the GDP state is the more stable state.

To further prove this assumption, one would have to determine the affinities for SEPT2:GTP-SEPT6:GTP and SEPT7:GDP-SEPT7:GTP dimers. However, suitable structures are not available in the PDB, neither with GTP nor with a GTP analogon.

Here, we have presented a MD workflow that allows for the calculation of inter-subunit binding free energies in septin complexes. Our approach is not limited to these specific septin dimers but may also be applied to other septin complexes provided the input structures are of sufficient quality. Our results indicate that GDP in the GTPase-active subunits SEPT2 and SEPT7 is a prominent factor that maintains G interface stability by coordination of the GDP ribose ring to conserved amino acids in the SUE. Removal of the nucleotide leads to a destabilization of the interface, concomitant with enhanced structural flexibility. Further research is required to validate if this is a common feature in all septin species or if it is a human septin-specific phenomenon.

## Supporting information

suppelmentary figures

## Acknowledgements

The authors acknowledge support by the state of Baden-Württemberg through bwHPC, the German Research Foundation (DFG) through grant INST 40/575-1 FUGG (JUSTUS 2 high performance computing cluster) and the staff of the Uni Ulm computer center for technical support.

## Author Contributions

BG designed and performed the simulations, analyzed the data and improved the manuscript. JL supported in setting up the simulations, analyzed the data and improved the manuscript. TG analyzed the data and wrote the manuscript.

## Supplementary items

**Video 1**. Full trajectory of a representative SEPT2:GDP-SEPT6:apo COM-pulling simulation.

**Video 2**. Full trajectory of a representative SEPT7:apo-SEPT7:apo COM-pulling simulation.

**Supplementary figures**

## Notes

### Competing Interest Statement

The authors have declared no competing interest.

## References

Abraham, Mark James, Teemu Murtola, Roland Schulz, Szilárd Páll, Jeremy C. Smith, Berk Hess, and Erik Lindah. 2015. “GROMACS: High Performance Molecular Simulations through Multi-Level Parallelism from Laptops to Supercomputers.” SoftwareX 1–2:19–25. doi: 10.1016/J.SOFTX.2015.06.001.

Bernetti, Mattia, and Giovanni Bussi. 2020. “Pressure Control Using Stochastic Cell Rescaling.” The Journal of Chemical Physics 153(11):114107. doi: 10.1063/5.0020514.

Bertin, Aurélie, Michael A. McMurray, Luong Thai, Galo Garcia, Violet Votin, Patricia Grob, Theresa Allyn, Jeremy Thorner, and Eva Nogales. 2010. “Phosphatidylinositol-4,5-Bisphosphate Promotes Budding Yeast Septin Filament Assembly and Organization.” Journal of Molecular Biology 404(4):711–31. doi: 10.1016/j.jmb.2010.10.002.

Brausemann, Anton, Stefan Gerhardt, Anne-Kathrin Schott, Oliver Einsle, Andreas Große-Berkenbusch, Nils Johnsson, and Thomas Gronemeyer. 2016. “Crystal Structure of Cdc11, a Septin Subunit from Saccharomyces Cerevisiae.” 193:157–61. doi: 10.1016/j.jsb.2016.01.004.

Brognara, Gabriel, Humberto D. Muni. Pereira, José Brandão-Neto, Ana Paula Ulian Araujo, and Richard Charles Garratt. 2019. “Revisiting SEPT7 and the Slippage of β-Strands in the Septin Family.” Journal of Structural Biology 207(1):67–73. doi: 10.1016/j.jsb.2019.04.015.

Bussi, Giovanni, Davide Donadio, and Michele Parrinello. 2007. “Canonical Sampling through Velocity Rescaling.” The Journal of Chemical Physics 126(1):014101. doi: 10.1063/1.2408420.

Castro, Danielle Karoline Silva do Vale, Sabrina Matos de Oliveira da Silva, Humberto D’Muniz Pereira, Joci Neuby Alves Macedo, Diego Antonio Leonardo, Napoleão Fonseca Valadares, Patricia Suemy Kumagai, José Brandão-Neto, Ana Paula Ulian Araújo, and Richard Charles Garratt. 2020. “A Complete Compendium of Crystal Structures for the Human SEPT3 Subgroup Reveals Functional Plasticity at a Specific Septin Interface.” IUCrJ 7(3):462–79. doi: 10.1107/S2052252520002973.

Cavini, Italo A., Diego A. Leonardo, Higor V.D. Rosa, Danielle K. S. V. Castro, Humberto D’Muniz Pereira, Napoleão F. Valadares, Ana P. U. Araujo, and Richard C. Garratt. 2021. “The Structural Biology of Septins and Their Filaments: An Update.” Frontiers in Cell and Developmental Biology 9. doi: 10.3389/FCELL.2021.765085.

Cool, Austin M., and Steffen Lindert. 2022. “Umbrella Sampling Simulations Measure Switch Peptide Binding and Hydrophobic Patch Opening Free Energies in Cardiac Troponin.” Journal of Chemical Information and Modeling. doi: 10.1021/ACS.JCIM.2C00508.

Darden, Tom, Darrin York, and Lee Pedersen. 1998. “Particle Mesh Ewald: An N⋅log(N) Method for Ewald Sums in Large Systems.” The Journal of Chemical Physics 98(12):10089. doi: 10.1063/1.464397.

Daura, Xavier, Bernhard Jaun, Dieter Seebach, Wilfred F. Van Gunsteren, and Alan E. Mark. 1998. “Reversible Peptide Folding in Solution by Molecular Dynamics Simulation.” Journal of Molecular Biology 280(5):925–32. doi: 10.1006/JMBI.1998.1885.

Durell, Stewart R., Bernard R. Brooks, and Arieh Ben-Naim. 1994. “Solvent-Induced Forces between Two Hydrophilic Groups.” Journal of Physical Chemistry 98(8):2198–2202. doi: 10.1021/J100059A038/ASSET/J100059A038.FP.PNG_V03.

Essmann, Ulrich, Lalith Perera, Max L. Berkowitz, Tom Darden, Hsing Lee, and Lee G. Pedersen. 1998. “A Smooth Particle Mesh Ewald Method.” The Journal of Chemical Physics 103(19):8577. doi: 10.1063/1.470117.

Farkasovsky, Marian, Peter Herter, Beate Voss, and Alfred Wittinghofer. 2005. “Nucleotide Binding and Filament Assembly of Recombinant Yeast Septin Complexes.” Biological Chemistry 386:643–56. doi: 10.1515/BC.2005.075.

Fischer, Martin, Dominik Frank, Reinhild Rösler, Nils Johnsson, and Thomas Gronemeyer. 2022. “Biochemical Characterization of a Human Septin Octamer.” Frontiers in Cell and Developmental Biology 10:771388. doi: 10.3389/FCELL.2022.771388.

Grupp, Benjamin, and Thomas Gronemeyer. 2022. “A Biochemical View on the Septins, a Less Known Component of the Cytoskeleton.” Biological Chemistry 0(0). doi: 10.1515/HSZ-2022-0263.

Hess, Berk. 2008. “P-LINCS: A Parallel Linear Constraint Solver for Molecular Simulation.” Journal of Chemical Theory and Computation 4(1):116–22. doi: 10.1021/CT700200B/ASSET/IMAGES/LARGE/CT700200BF00001.JPEG.

Hess, Berk, Henk Bekker, Herman J. C. Berendsen, and Johannes G. E. M. Fraaije. 1997. “LINCS: A Linear Constraint Solver for Molecular Simulations.” J Comput Chem 18:14631472. doi: 10.1002/(SICI)1096-987X(199709)18:12.

Huang, Jing, Sarah Rauscher, Grzegorz Nawrocki, Ting Ran, Michael Feig, Bert L. De Groot, Helmut Grubmüller, and Alexander D. MacKerell. 2017. “CHARMM36m: An Improved Force Field for Folded and Intrinsically Disordered Proteins.” Nature Methods 14(1):71–73. doi: 10.1038/NMETH.4067.

Hub, Jochen S., Bert L. De Groot, and David Van Der Spoel. 2010. “G-Whams-a Free Weighted Histogram Analysis Implementation Including Robust Error and Autocorrelation Estimates.” Journal of Chemical Theory and Computation 6(12):3713–20. doi: 10.1021/ct100494z.

Isralewitz, Barry, Jerome Baudry, Justin Gullingsrud, Dorina Kosztin, and Klaus Schulten. 2001. “Steered Molecular Dynamics Investigations of Protein Function.” Journal of Molecular Graphics and Modelling 19(1):13–25. doi: 10.1016/S1093-3263(00)00133-9.

Jorgensen, William L., Jayaraman Chandrasekhar, Jeffry D. Madura, Roger W. Impey, and Michael L. Klein. 1998. “Comparison of Simple Potential Functions for Simulating Liquid Water.” The Journal of Chemical Physics 79(2):926. doi: 10.1063/1.445869.

Kästner, Johannes. 2011. “Umbrella Sampling.” Wiley Interdisciplinary Reviews: Computational Molecular Science 1(6):932–42. doi: 10.1002/WCMS.66.

Kinoshita, Makoto. 2003. “The Septins.” Genome Biology 4(11):236. doi: 10.1186/gb-2003-4-11-236.

Klepeis, John L., Kresten Lindorff-Larsen, Ron O. Dror, and David E. Shaw. 2009. “Long-Timescale Molecular Dynamics Simulations of Protein Structure and Function.” Current Opinion in Structural Biology 19(2):120–27. doi: 10.1016/J.SBI.2009.03.004.

Kumar, Shankar, John M. Rosenberg, Djamal Bouzida, Robert H. Swendsen, and Peter A. Kollman. 1992. “The Weighted Histogram Analysis Method for Free-Energy Calculations on Biomolecules. I. The Method.” Journal of Computational Chemistry 13(8):1011–21. doi: 10.1002/JCC.540130812.

Lee, Jumin, Xi Cheng, Jason M. Swails, Min Sun Yeom, Peter K. Eastman, Justin A. Lemkul, Shuai Wei, Joshua Buckner, Jong Cheol Jeong, Yifei Qi, Sunhwan Jo, Vijay S. Pande, David A. Case, Charles L. Brooks, Alexander D. MacKerell, Jeffery B. Klauda, and Wonpil Im. 2016. “CHARMM-GUI Input Generator for NAMD, GROMACS, AMBER, OpenMM, and CHARMM/OpenMM Simulations Using the CHARMM36 Additive Force Field.” Journal of Chemical Theory and Computation 12(1):405–13. doi: 10.1021/ACS.JCTC.5B00935/ASSET/IMAGES/LARGE/CT-2015-00935E_0005.JPEG.

Lemkul, Justin A. 2019. “From Proteins to Perturbed Hamiltonians: A Suite of Tutorials for the GROMACS-2018 Molecular Simulation Package [Article v1.0].” Living Journal of Computational Molecular Science 1(1):5068–5068. doi: 10.33011/LIVECOMS.1.1.5068.

Lemkul, Justin A., and David R. Bevan. 2010. “Assessing the Stability of Alzheimer’s Amyloid Protofibrils Using Molecular Dynamics.” Journal of Physical Chemistry B 114(4):1652–60. doi: 10.1021/jp9110794.

Lu, Chao, Chuanjie Wu, Delaram Ghoreishi, Wei Chen, Lingle Wang, Wolfgang Damm, Gregory A. Ross, Markus K. Dahlgren, Ellery Russell, Christopher D. Von Bargen, Robert Abel, Richard A. Friesner, and Edward D. Harder. 2021. “OPLS4: Improving Force Field Accuracy on Challenging Regimes of Chemical Space.” Journal of Chemical Theory and Computation 17(7):4291–4300. doi: 10.1021/ACS.JCTC.1C00302.

Macedo, Joci N. A., Napoleão F. Valadares, Ivo A. Marques, Frederico M. Ferreira, Julio C. P. Damalio, Humberto M. Pereira, Richard C. Garratt, and Ana P. U. Araujo. 2013. “The Structure and Properties of Septin 3: A Possible Missing Link in Septin Filament Formation.” The Biochemical Journal 450(1):95–105. doi: 10.1042/BJ20120851.

Madhavi Sastry, G., Matvey Adzhigirey, Tyler Day, Ramakrishna Annabhimoju, and Woody Sherman. 2013. “Protein and Ligand Preparation: Parameters, Protocols, and Influence on Virtual Screening Enrichments.” Journal of Computer-Aided Molecular Design 27(3):221–34. doi: 10.1007/S10822-013-9644-8.

Mendonça Deborah C., Samuel L. Guimarães, Humberto D’Muniz Pereira, Andressa A. Pinto, Marcelo A. de Farias, Andre S. de Godoy, Ana P. U. Araujo, Marin van Heel, Rodrigo V. Portugal, and Richard C. Garratt. 2021. “An Atomic Model for the Human Septin Hexamer by Cryo-EM.” Journal of Molecular Biology 433(15):167096. doi: 10.1016/j.jmb.2021.167096.

Mendonça Deborah C., Joci N. Macedo, Samuel L. Guimarães, Fernando L. Barroso da Silva, Alexandre Cassago, Richard C. Garratt, Rodrigo V. Portugal, and Ana P. U. Araujo. 2019. “A Revised Order of Subunits in Mammalian Septin Complexes.” Cytoskeleton 76(9– 10):457–66. doi: 10.1002/cm.21569.

Menon, Manoj B. 2018. “Septin.” Pp. 4875–84 in Encyclopedia of Signaling Molecules. Cham: Springer International Publishing.

Mostowy, Serge, and Pascale Cossart. 2012. “Septins: The Fourth Component of the Cytoskeleton.” Nature Reviews. Molecular Cell Biology 13(3):183–94. doi: 10.1038/nrm3284.

Neria, Eyal, Stefan Fischer, and Martin Karplus. 1998. “Simulation of Activation Free Energies in Molecular Systems.” The Journal of Chemical Physics 105(5):1902. doi: 10.1063/1.472061.

Pan, Fangfang, Russell L. Malmberg, and Michelle Momany. 2007. “Analysis of Septins across Kingdoms Reveals Orthology and New Motifs.” BMC Evol Biol 7:103. doi: 10.1186/1471-2148-7-103.

Parrinello, M., and A. Rahman. 1998. “Polymorphic Transitions in Single Crystals: A New Molecular Dynamics Method.” Journal of Applied Physics 52(12):7182. doi: 10.1063/1.328693.

Patel, Jagdish Suresh, and F. Marty Ytreberg. 2018. “Fast Calculation of Protein-Protein Binding Free Energies Using Umbrella Sampling with a Coarse-Grained Model.” Journal of Chemical Theory and Computation 14(2):991–97. doi: 10.1021/ACS.JCTC.7B00660.

Peterson, EA, and EM Petty. 2010. “Conquering the Complex World of Human Septins: Implications for Health and Disease.” Clinical Genetics 77(6):511–24. doi: 10.1111/j.1399-0004.2010.01392.x.

Rosa, Higor Vinícius Dias, Diego Antonio Leonardo, Gabriel Brognara, José Brandão-Neto, Humberto D’Muniz Pereira, Ana Paula Ulian Araújo, and Richard Charles Garratt. 2020. “Molecular Recognition at Septin Interfaces: The Switches Hold the Key.” Journal of Molecular Biology 432(21):5784–5801. doi: 10.1016/j.jmb.2020.09.001.

Sheffield, Peter J., Carey J. Oliver, Brandon E. Kremer, Sitong Sheng, Zhifeng Shao, and Ian G. Macara. 2003. “Borg/Septin Interactions and the Assembly of Mammalian Septin Heterodimers, Trimers, and Filaments.” The Journal of Biological Chemistry 278(5):3483–88. doi: 10.1074/jbc.M209701200.

Shuman, Brent, and Michelle Momany. 2022. “Septins From Protists to People.” Frontiers in Cell and Developmental Biology 9:824850. doi: 10.3389/FCELL.2021.824850.

Sirajuddin, Minhajuddin, Marian Farkasovsky, Florian Hauer, Dorothee Kühlmann, Ian G. Macara, Michael Weyand, Holger Stark, and Alfred Wittinghofer. 2007. “Structural Insight into Filament Formation by Mammalian Septins.” Nature 449(7160):311–15. doi: 10.1038/nature06052.

Valadares, Napoleão Fonseca, Humberto d’ Muniz Pereira, Ana Paula Ulian Araujo, and Richard Charles Garratt. 2017. “Septin Structure and Filament Assembly.” Biophysical Reviews 9(5):481–500. doi: 10.1007/S12551-017-0320-4/FIGURES/12.

Versele, Matthias, and Jeremy Thorner. 2004. “Septin Collar Formation in Budding Yeast Requires GTP Binding and Direct Phosphorylation by the PAK, Cla4.” Journal of Cell Biology 164(5):701–15. doi: 10.1083/jcb.200312070.

Vrabioiu, Alina M., Scott A. Gerber, Steven P. Gygi, Christine M. Field, and Timothy J. Mitchison. 2004. “The Majority of the Saccharomyces Cerevisiae Septin Complexes Do Not Exchange Guanine Nucleotides.” Journal of Biological Chemistry 279(4):3111–18. doi: 10.1074/jbc.M310941200.

Weems, Andrew, and Michael McMurray. 2017. “The Step-Wise Pathway of Septin Hetero-Octamer Assembly in Budding Yeast.” ELife 6:e23689. doi: 10.7554/eLife.23689.

Weirich, Christine S., Jan P. Erzberger, and Yves Barral. 2008. “The Septin Family of GTPases: Architecture and Dynamics.” Nature Reviews. Molecular Cell Biology 9(6):478–89. doi: 10.1038/nrm2407.

Zent, Eldar, Ingrid Vetter, and Alfred Wittinghofer. 2011. “Structural and Biochemical Properties of Sept7, a Unique Septin Required for Filament Formation.” Biological Chemistry 392:791–97. doi: 10.1515/BC.2011.082.

Zent, Eldar, and Alfred Wittinghofer. 2014. “Human Septin Isoforms and the GDP-GTP Cycle.” Biological Chemistry 395(2):169–80. doi: 10.1515/HSZ-2013-0268/MACHINEREADABLECITATION/RIS.

Zheng, Liangzhen, Amr A. Alhossary, Chee Keong Kwoh, and Yuguang Mu. 2018. “Molecular Dynamics and Simulation.” Encyclopedia of Bioinformatics and Computational Biology: ABC of Bioinformatics 1–3(V):550–66. doi: 10.1016/B978-0-12-809633-8.20284-7.

