## Supplementary material for "An *in silico* approach to determine inter-subunit affinities in human septin complexes": suppelmentary figures

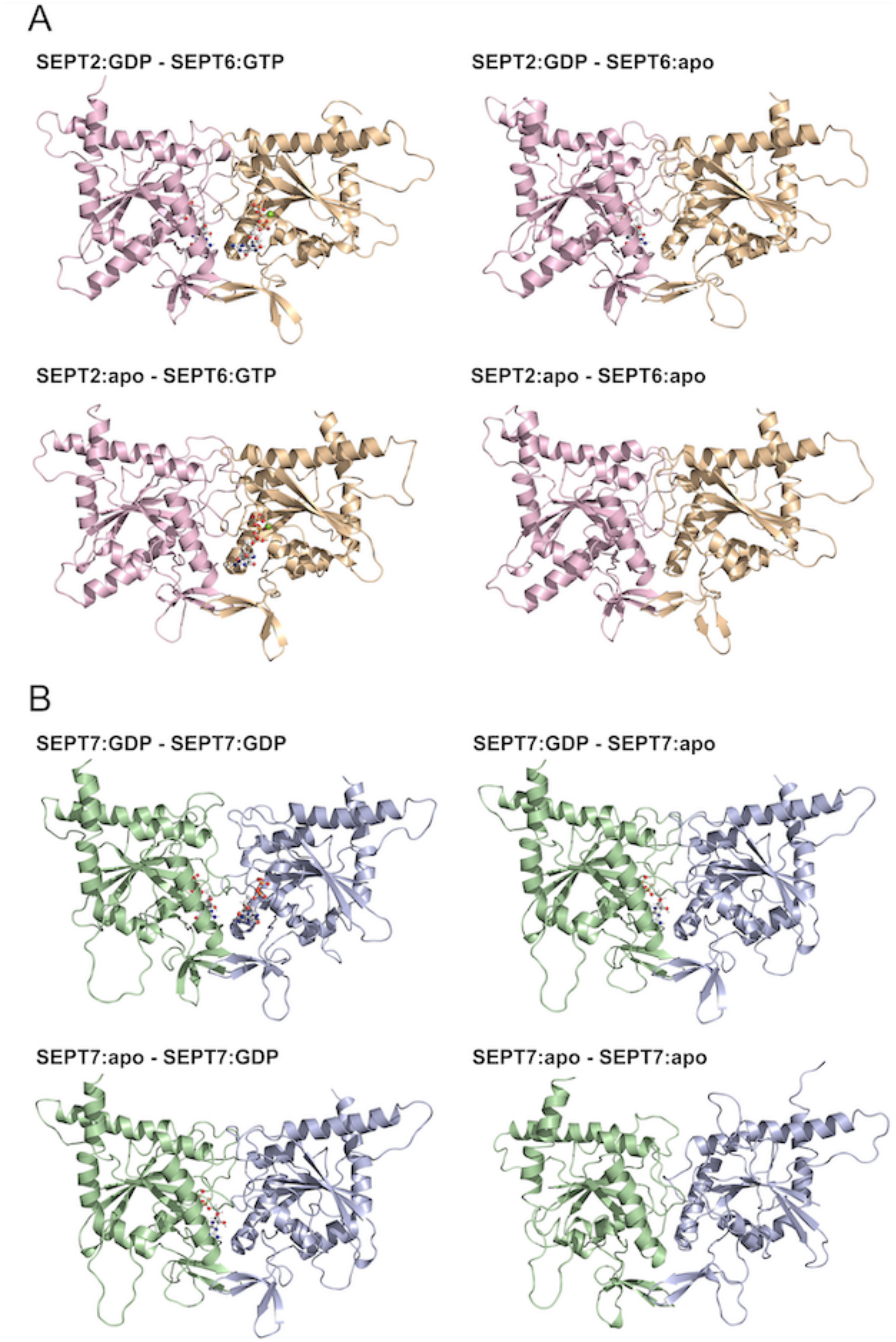


**Suppl. Fig. 1.** Relaxed input structures for the COM-pulling simulations of all employed septin dimers. **(A)** SEPT-SEPT6 dimers, **(B**) SEPT7-SEPT7 dimers.

SEPT7 subunits are colored palegreen and lightblue, SEPT2 lightpink and SEPT6 wheat.

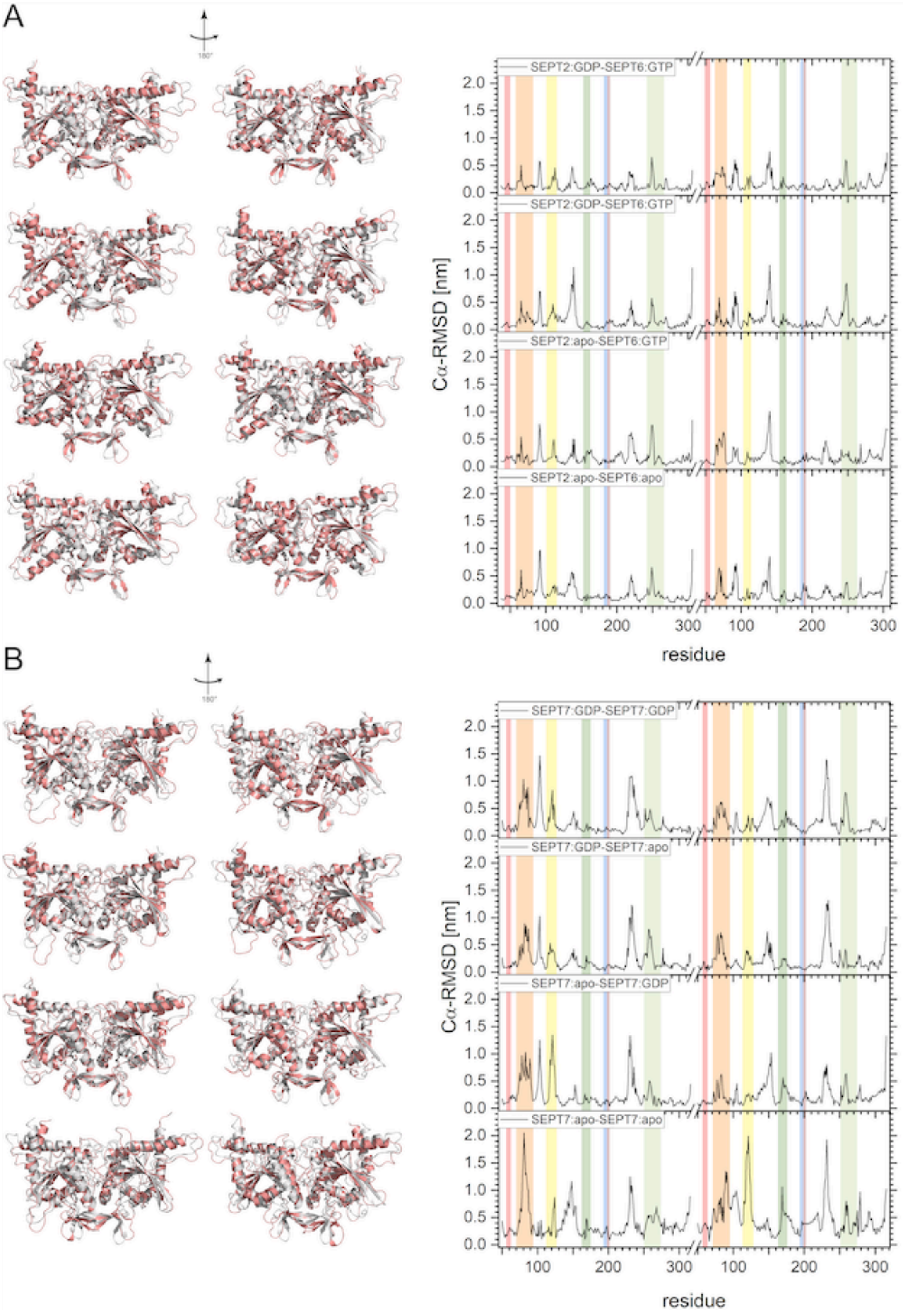


**Suppl. Fig. 2.** Cα-RMSD of the relaxed structures referenced to the input structures for **(A)** all SEPT2-SEPT6 dimers and **(B)** all SEPT7-SEPT7 dimers.

An overlay (left) of the relaxed structures (salmon) with the input structures (grey) and the corresponding RMSD plots (right) are shown. The color coding in the RMSD plots corresponds to the structural elements highlighted in Fig. 1.

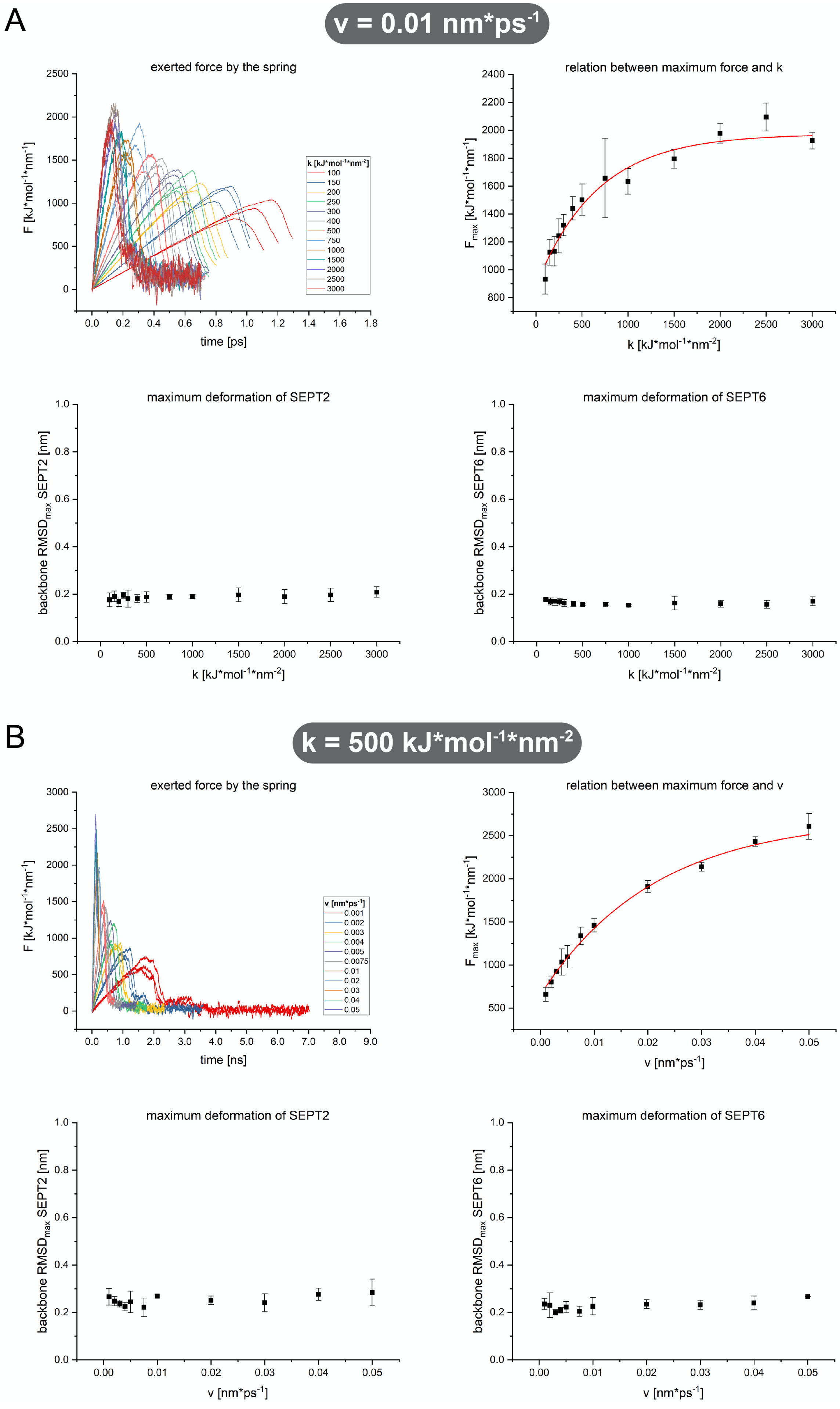
A
v = 0.01 nm*ps-1
exerted force by the spring
F [kJ*mol-1*nm-1]
time [ps]
k [kJ*mol-1*nm-2]
100
150
200
250
300
400
500
750
1000
1500
2000
2500
3000
relation between maximum force and k
Fmax [kJ*mol-1*nm-1]
k [kJ*mol-1*nm-2]
maximum deformation of SEPT2
backbone RMSDmax SEPT2 [nm]
k [kJ*mol-1*nm-2]
maximum deformation of SEPT6
backbone RMSDmax SEPT6 [nm]
k [kJ*mol-1*nm-2]
B
k = 500 kJ*mol-1*nm-2
exerted force by the spring
F [kJ*mol-1*nm-1]
time [ns]
v [nm*ps-1]
0.001
0.002
0.003
0.004
0.005
0.0075
0.01
0.02
0.03
0.04
0.05
relation between maximum force and v
Fmax [kJ*mol-1*nm-1]
v [nm*ps-1]
maximum deformation of SEPT2
backbone RMSDmax SEPT2 [nm]
v [nm*ps-1]
maximum deformation of SEPT6
backbone RMSDmax SEPT6 [nm]
v [nm*ps-1]

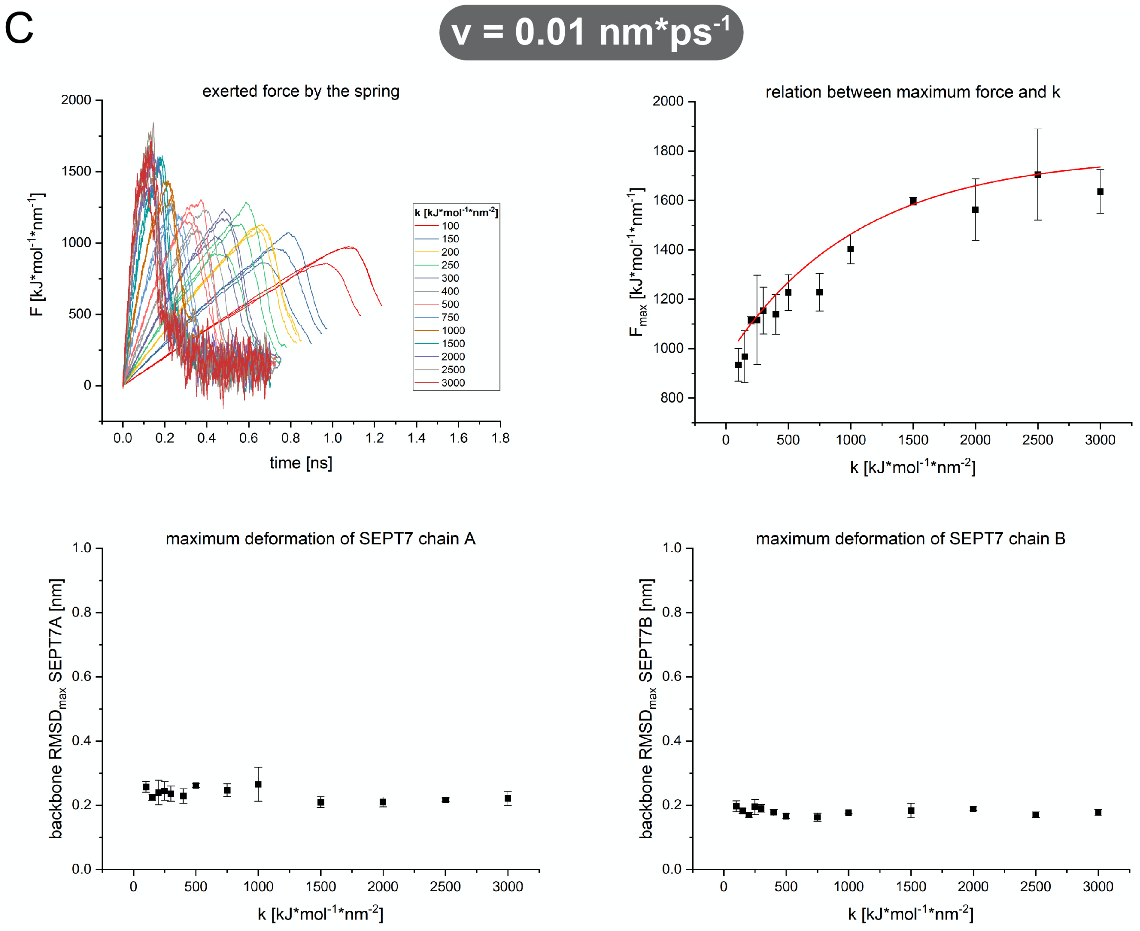

C
v = 0.01 nm*ps-1
exerted force by the spring
F [kJ*mol-1*nm-1]
time [ns]
k [kJ*mol-1*nm-2]
100
150
200
250
300
400
500
750
1000
1500
2000
2500
3000
relation between maximum force and k
Fmax [kJ*mol-1*nm-1]
k [kJ*mol-1*nm-2]
maximum deformation of SEPT7 chain A
backbone RMSDmax SEPT7A [nm]
k [kJ*mol-1*nm-2]
maximum deformation of SEPT7 chain B
backbone RMSDmax SEPT7B [nm]
k [kJ*mol-1*nm-2]


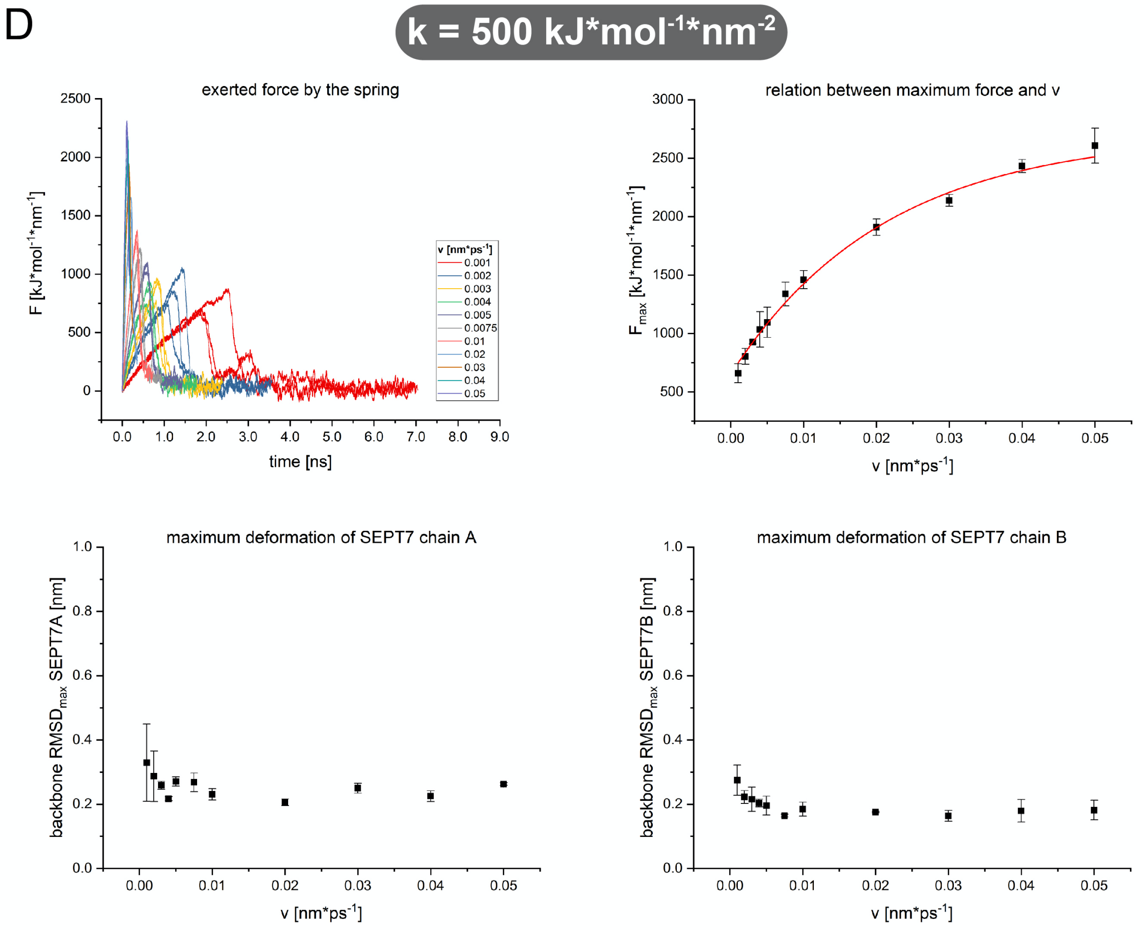

D
k = 500 kJ*mol-1*nm-2
exerted force by the spring
F [kJ*mol-1*nm-1]
time [ns]
v [nm*ps-1]
0.001
0.002
0.003
0.004
0.005
0.0075
0.01
0.02
0.03
0.04
0.05
relation between maximum force and v
Fmax [kJ*mol-1*nm-1]
v [nm*ps-1]
maximum deformation of SEPT7 chain A
backbone RMSDmax SEPT7A [nm]
v [nm*ps-1]
maximum deformation of SEPT7 chain B
backbone RMSDmax SEPT7B [nm]
v [nm*ps-1]

**Suppl. Fig. 3.** Analysis of different combinations of k and v in the COM-pulling simulations with the dimers SEPT2:GDP-SEPT6:GTP **(A), (B)** and SEPT7:GDP-SEPT7:GDP **(C), (D)**. In (A) and (C) v was kept constant at 0.01 $nm \cdot ps^{-1}$ and k was varied from 100 – 3000 $kJ \cdot mol^{-1} \cdot$

$nm^{-2}$. In (B) and (D) k was kept constant at 500 $kJ \cdot mol^{-1} \cdot nm^{-2}$ and v was varied from 0.001 – 0.05 $nm \cdot ps^{-1}$. The upper-left graph in each panel shows the force exerted by the spring. The upper-right graph in each panel shows the behavior of the maximum observed force for the given combination of v and k. In the lower graphs, the maximum structural deformation of the different subunits (backbone $RMSD_{max}$) is shown which was calculated by RMSD-based alignments of the backbone atoms (N, Ca, C) in the respective subunit. The latter analysis was performed to identify possible unfolding artifacts introduced during the COM-pulling simulations. All simulations were performed in triplicate. Error bars represent the standard deviation.

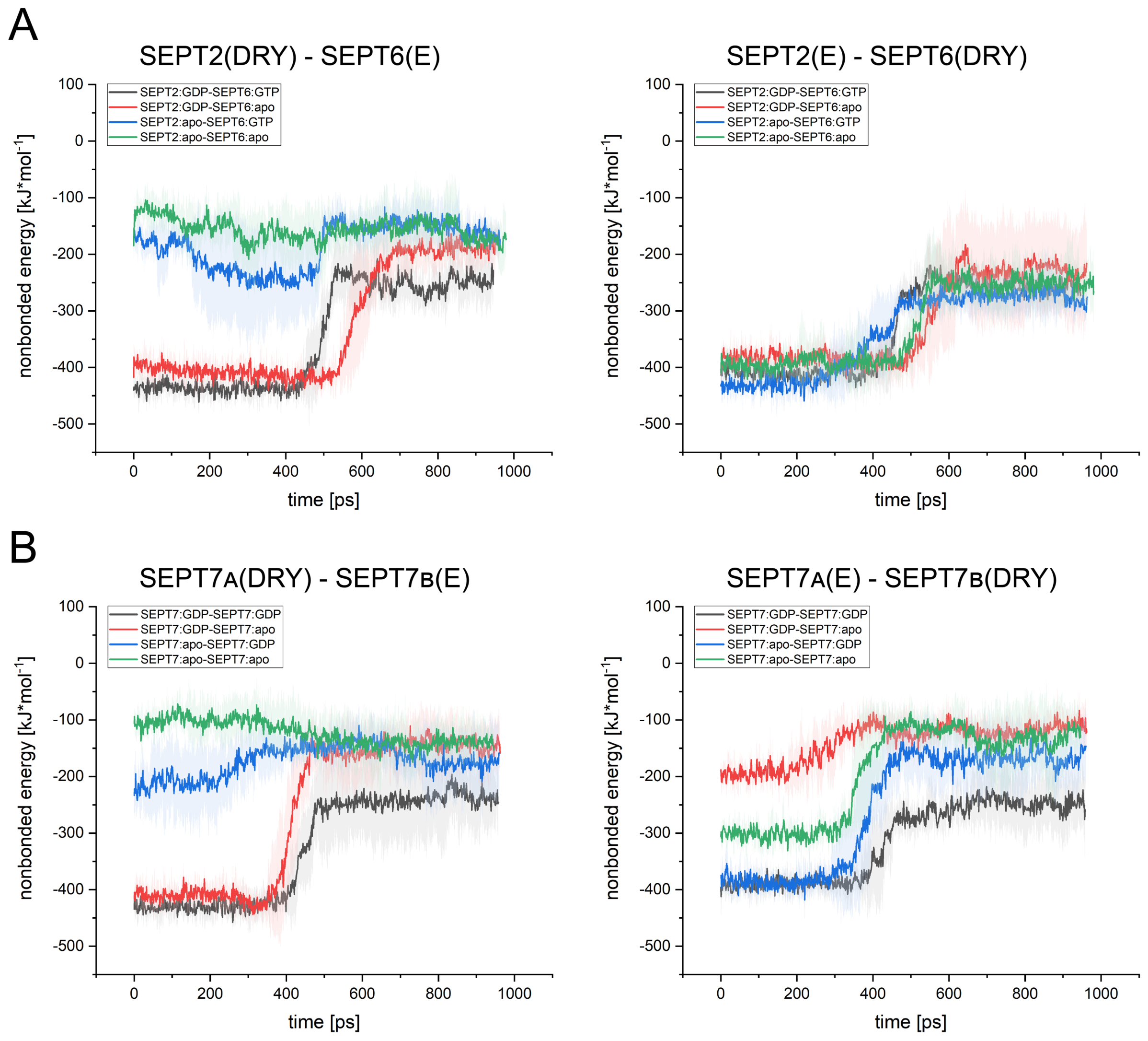


**Suppl. Fig. 4**. Nonbonded energies plotted vs. the simulation time in SEPT2-6 and SEPT7-7 dimers. **(A)** Nonbonded energies of all nucleotide binding states for the binding pocket in SEPT2 (left) and SEPT6 (right) in the SEPT2-SEPT6 dimer. Note that the energies remain unchanged upon nucleotide removal in SEPT6. **(B)** Nonbonded energies of all nucleotide binding states for the binding pocket in SEPT7 chain A (left) and SEPT7 chain B (right) in the SEPT7 dimer.

DRY abbreviate Asp(G4), Arg(βb), Tyr(βb) in the binding pocket and E abbreviates Glu(Tr2) from the neighboring subunit, respectively. Error bands represent the standard deviation.
